# OmicsFootPrint: a framework to integrate and interpret multi-omics data using circular images and deep neural networks

**DOI:** 10.1101/2024.03.21.586001

**Authors:** Xiaojia Tang, Naresh Prodduturi, Kevin J. Thompson, Richard Weinshilboum, Ciara C. O’Sullivan, Judy C. Boughey, Hamid R. Tizhoosh, Eric W. Klee, Liewei Wang, Matthew P. Goetz, Vera Suman, Krishna R. Kalari

## Abstract

The OmicsFootPrint framework addresses the need for advanced multi-omics data analysis methodologies by transforming data into intuitive two-dimensional circular images and facilitating the interpretation of complex diseases. Utilizing Deep Neural Networks and incorporating the SHapley Additive exPlanations (SHAP) algorithm, the framework enhances model interpretability. Tested with The Cancer Genome Atlas (TCGA) data, OmicsFootPrint effectively classified lung and breast cancer subtypes, achieving high Area Under Curve (AUC) scores— 0.98±0.02 for lung cancer subtype differentiation, 0.83±0.07 for breast cancer PAM50 subtypes, and successfully distinguished between invasive lobular and ductal carcinomas in breast cancer, showcasing its robustness. It also demonstrated notable performance in predicting drug responses in cancer cell lines, with a median AUC of 0.74, surpassing nine existing methods. Furthermore, its effectiveness persists even with reduced training sample sizes. OmicsFootPrint marks an enhancement in multi-omics research, offering a novel, efficient, and interpretable approach that contributes to a deeper understanding of disease mechanisms.

## INTRODUCTION

Multi-omics data, including genomics, transcriptomics, epigenomics, proteomics, and metabolomics, offer diverse molecular perspectives of individual biological samples. These high-dimensional data sets are pivotal in revealing hidden biological information and facilitating the discovery of pathophysiologic mechanisms underlying disease (1–4). The advent of deep learning technologies has revolutionized the analysis of multi-omics data, particularly in complex disorders such as cancer and neurodegenerative diseases. Deep Neural Networks (DNNs) and Graph Neural Networks (GNNs), with their ability to capture complex patterns across multiple layers of information, have proven instrumental in identifying novel biomarkers, deciphering cancer progression, and predicting responses to therapies(5–8). This has opened new avenues for understanding tumor biology and has been a catalyst in the movement toward personalized and precise cancer treatments(9–11).

Despite progress in omics data science, progress in the development of effective methodologies to combine different types of omics data has been slow. However, combining these data types is essential for a deeper understanding of biological systems. It will result in more precise disease models, tailored treatments in personalized medicine, and richer insight into complex biological mechanisms. As the field rapidly advances, deep learning is mainly being used to analyze structured, table-like multi-omics data. Several strategic multi-omics integration methods have been developed, including early integration (combining multi-omics data at analysis onset), late integration (combining multi-omics data at the end of analyses), and hybrid approaches (using both early and late integration techniques). Early integration techniques focus on merging multiple omics data sets to decipher an integrated latent structure.

A notable example of early integration is the pioneering work by Chaudhary et al. (7), who employed an autoencoder-based deep learning framework for a multi-omics hepatocellular carcinoma (HCC) data set. Similarly, Xie et al. made strides with the GDP (Group lasso regularized Deep learning for cancer Prognosis) algorithm, combining a deep learning framework based on TensorFlow (Google Inc.) with the Cox proportional hazards model for multi-omics driven cancer survival analysis (12). Conversely, late integration methods independently extract latent feature vectors from each omics type, subsequently amalgamating them into a cohesive representation. This is exemplified by the MOLI (Multi-Omics Late Integration) method, which extracts features from gene expression, copy number, and somatic mutation data and then merges them using a dual cost function involving triplet loss and binary cross-entropy loss for enhanced optimization (13). Simidjievski et al. implemented autoencoder architectures (14), while Wang et al. incorporated a graphical convolution network in their MOGONET design (15). Benkirane et al. developed a mixed integration approach that combined the advantage of both early and late integrations, also based on autoencoder methodologies(16). However, it is critical to note that the efficacy of deep learning in this domain is significantly dependent on the availability of large training data sets. This requirement poses a notable challenge for high-throughput multi-omics data, which is characteristically high-dimensional and plagued by low sample sizes (HDLSS), often leading to issues of model overfitting (17,18).

One of the solutions to the HDLSS challenge in high-throughput multi-omics data is the transformation of tabular data into image format, leveraging the success of DNNs in image analysis. Methods like DeepInsight (19), REFINED (20), and IGTD (21) have been developed to analyze omics features from a single platform (mainly gene expression) into 2-D spaces based on pairwise similarities (19–21). However, these methods have limitations. They are susceptible to biases and overfitting, especially when the distribution of samples is skewed. Additionally, they are currently limited to analyzing data from only one type of omics data set. A recent method, DeepInsight-3D, represents an innovative step forward in the realm of high-throughput multi-omics data analysis(22). Building upon its predecessor, DeepInsight extends its capabilities to handle multi-omics data more effectively. Unlike traditional approaches that are limited to processing a single type of omics data, DeepInsight-3D is designed to integrate up to three different omics types concurrently. This is achieved by employing a sophisticated framework that allows for the co-localization of features from diverse omics types, such as gene expression, proteomics, or metabolomics, in a three-dimensional space. However, DeepInsight-3D still grapples with certain limitations. The positioning of features from the other two omics types is heavily dependent on the spatial arrangement of the gene expression data. This dependence can introduce intrinsic biases, mainly when the feature distribution in the gene expression is uneven, potentially undermining the utilization of the other two omics features. Given that the gene expression data dominates the spatial arrangement in DeepInsight-3D, the nuances and distinct characteristics of the other two omics data may not be adequately represented or integrated, thus diminishing the holistic insight that multi-omics integration aims to provide. Additionally, DeepInsight-3D’s capacity to integrate only three types of omics data can be a constraint, especially considering the vast and varied nature of multi-omics data sets available in contemporary research.

To overcome these challenges, we developed OmicsFootPrint, a novel framework in the field of multi-omics data analysis. This method organizes multi-omics features into a circular plot based on chromosomal locations (23). This organization aims to facilitate deep learning algorithms in discerning relationships among spatially associated multi-omics features to enhance the accuracy of image classification and the correlation of molecular characteristics with various phenotypes. Additionally, OmicsFootPrint incorporates the Shapley Additive exPlanations (SHAP) algorithm for interpreting the trained models, providing insight into the data driving the model results and identifying complex patterns that traditional methods might miss. In this study, we evaluated various deep-learning architectures and multi-omics data combinations to identify the most effective approaches using The Cancer Genome Atlases (TCGA) lung and breast cancer data sets. We benchmarked our method with a publicly available cancer cell lines drug response data set and demonstrated OmicsFootPrint’s superiority over existing algorithms. We further applied OmicsFootPrint for histological subtyping of TCGA breast cancer tumors, utilizing four types of omics data: RNA-seq gene expression, copy number variation (CNV), protein assay, and microRNA, showcasing our framework findings, interpretability of models, versatility, and robustness in multi-omics data analysis.

## Materials and METHODS

### OmicsFootPrint Framework

The OmicsFootPrint framework is composed of three key modules: image generation and preprocessing (stage 1), convolutional neural network (CNN) training and prediction (stage 2), and decoding using SHAP analysis (stage 3), as shown in Figure 1. A detailed description of the modules is presented below.

**Figure 1.**
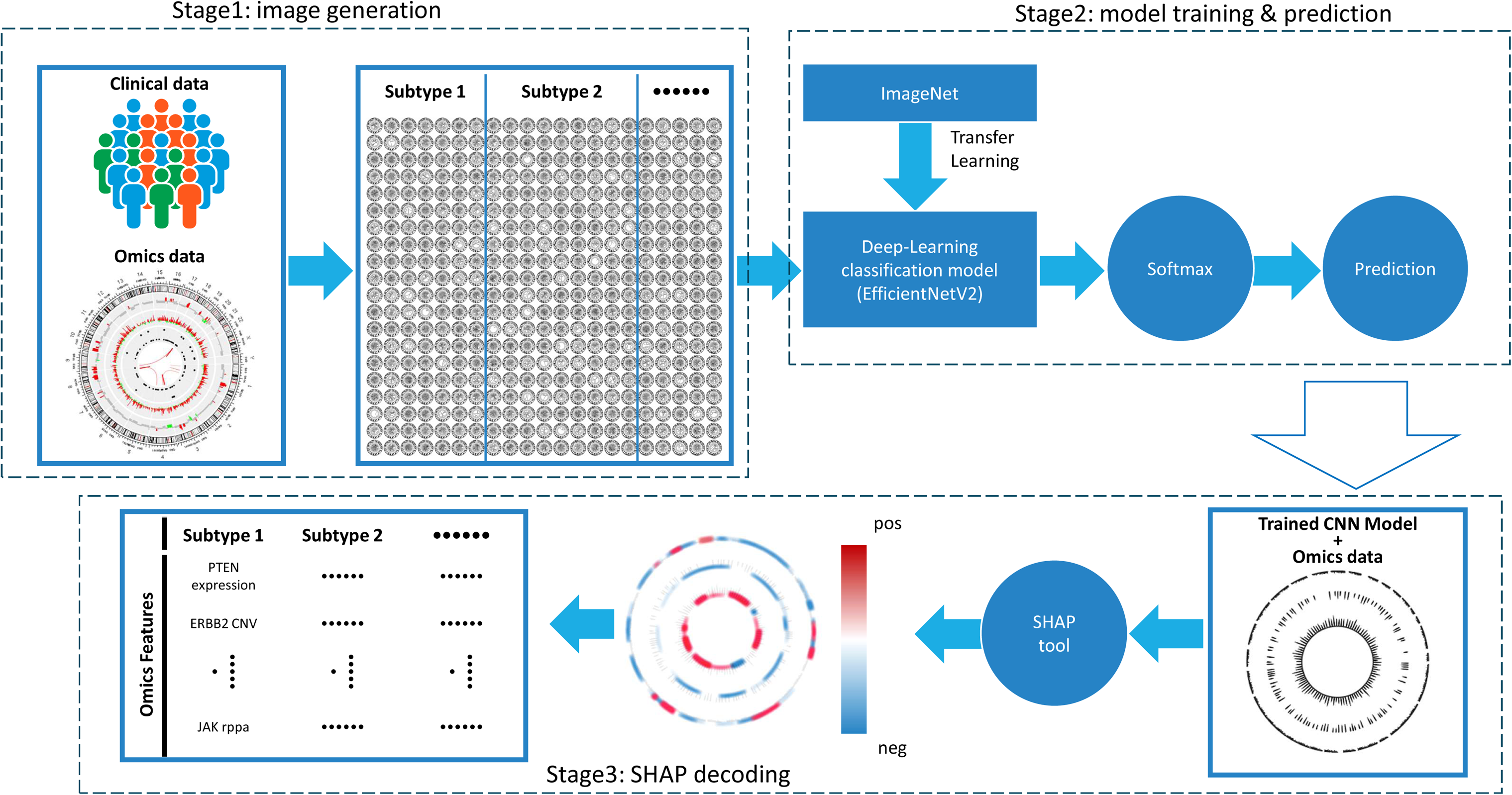
A high-level overview of the OmicsFootPrint workflow. CNN indicates convolutional neural network; CNV, copy number variation; neg, negative; pos, positive; SHAP, SHapley Additive exPlanations.

### Image generation

Multi-omics data for each individual is transformed into circular images(23), where each omics data type is represented in a separate circular track based on the genomic locations (i.e., the chromosome and position along the chromosome). In each track, the features are arranged based on their genomic location along the circle, from chromosomes 1-22. Based on the biological questions and data availability, the sex or mitochondrial chromosomes may be optional. Users can customize the layout arrangement of the data type on each track. By default, omics data types with more features are assigned to outer tracks, and those with fewer features to inner tracks. The height of each track was standardized across all samples, determined by the range of values for each feature. This method ensured consistency in the representation of data across different samples. The resulting circular images were initially constructed as 1024×1024 pixels. Figure 2 shows an example that includes the circular diagram layout of an individual patient from TCGA with copy number data, Reverse Phase Protein Array (RPPA), and expression (from outer to inner track, respectively), as well as its OmicsFootPrint black and white image that is used in the training of deep-learning models.

**Figure 2.**
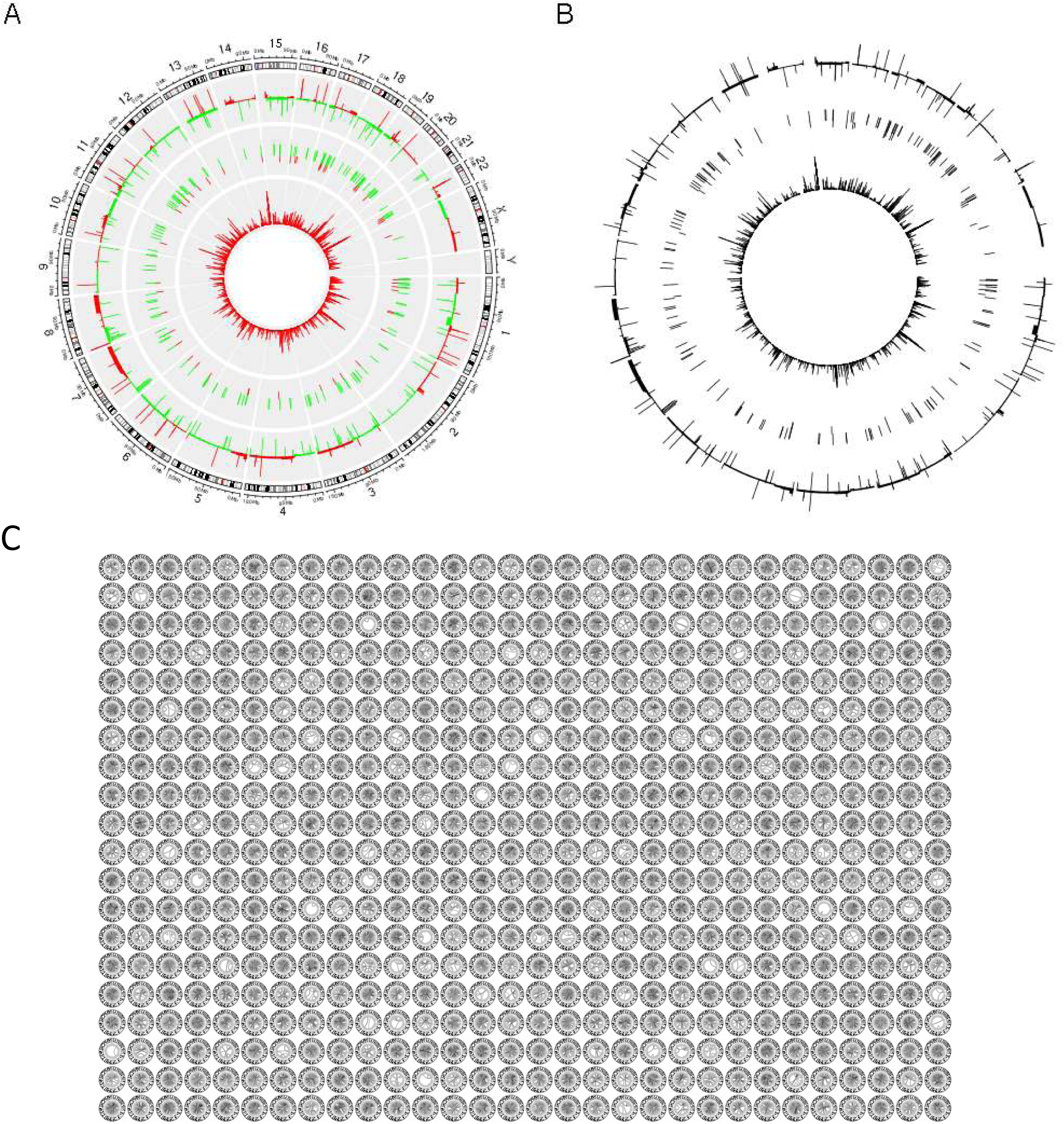
The illustration of the converted circular images (A) shows a detailed circular image with multiple tracks. Each track represents different omics data types, such as gene expression levels, copy number variations, or other genomic measurements. (B) It presents a simplified circular image that was used as input for the OmicsFootPrint framework. The data points are uniformly distributed along the circumference, which represents a single type of omics data or a binary representation of the presence or absence of certain features. (C) Displays OmicsFootPrints of 700 samples in a grid format. This represents a high-throughput analysis of circular plots corresponding to individual samples or experimental conditions. Patterns across multiple samples are identified using these images.

### Image preprocessing and data set preparation

The input circular footprint images (1024×1024 pixels) were rescaled internally to 256×256, and feature scaling of the input image data was done by dividing by 255. By default, the OmicsFootPrint image set was randomly divided into three subsets: training (70%), testing (15%), and validation (15%). Ten permutations, each with a different random seed, were performed to enhance the robustness and generalizability of the performance assessment.

### Model training

To address the challenge of relatively small sample sizes in the clinical studies, we initialized the deep-learning models pre-with ImageNet weights and applied transfer learning (24). We also used several image augmentation techniques to enhance the diversity of the training data sets, including random flipping of images (both horizontally and vertically) and introducing random variability in image brightness, saturation, hue, and contrast using the TensorFlow python package with a 50% randomness. Such augmentation techniques are crucial for enhancing the robustness and generalization of the model training.

A SoftMax function was used to activate the output layer. The training process is performed by minimizing the cross-entropy loss function. Considering memory constraints, we set the training batch size at 32 samples and ran the models for 100 epochs, with an evaluation taking place during each epoch. However, the batch size should be optimized based on the available computational resources. We selected the Nadam optimizer, initiating it with starting a learning rate of 1 × 10^−5^; this approach applied categorical cross-entropy for multi-class classification and binary cross-entropy for binary classification (25). To address class imbalance, class weights were adjusted to be inversely proportional to the class frequencies. Further optimization of hyperparameters such as dropout, optimizer, and learning rate was performed using grid searches.

### Model performance evaluation

The performance of the model was evaluated using an area under the curve (AUC) metric. The AUC was calculated for an independent test set.

### Model selection

The OmicsFootPrint framework is designed to be flexible, allowing for the integration of various deep-learning models for image classification. In this study, we compared several state-of-the-art deep neural network (DNN) architectures, including VGG16 combined with AutoGluon (26,27), bilinear CNN (BCNN) (28), DenseNet-121 (29), and EfficientNetV2 (30).

The EfficientNetV2-Large (30) and DenseNet-121 (29) architectures were initialized, with default weights transferred from the ImageNet data set. In this setup, all layers were frozen except for the final fully connected layer. The output tensor of the base model was processed through a series of layers: 1) a global average pooling 2D layer to reduce spatial dimensions (31); 2) a flattened layer to convert the 2D matrix into a 1D vector; 3) a dense layer with 512 units where ReLU activation was employed; and 4) a final output layer with a SoftMax activation function designed to predict subtypes was implemented.

Both VGG+AutoGluon and BCNN models were built on the VGG16 architecture, which was initialized with default weights transferred from the ImageNet data set. In the VGG+AutoGluon model, we removed the output layer, yielding a final fully connected layer with 4,096 output nodes. This modification enabled the formation of a feature vector output for the images. The images were then preprocessed, and the resulting model was used to extract the feature vectors. Dimension reduction was then performed on the most relevant features of the vectors using Principal Component Analysis (26). Finally, AutoGluon’s automated machine learning was used to train the model and predict the specific subtypes (27).

For the BCNN model (28), the "block4_conv3" convolutional layer was used as the starting point. Two dropout layers were added to this layer, followed by a custom lambda function, which calculated the outer product, implementing a bilinear pooling technique to capture pairwise feature interactions. The resulting bilinear tensor was then flattened into a 1D vector, with a subsequent dropout layer applied for regularization. This processed tensor was then passed through a dense layer consisting of 512 units with ReLU activation, followed by a final output layer equipped with a SoftMax activation function used for predictions. Regularization strategies were applied in the dense layers, utilizing both kernel and bias L2 regularization to prevent overfitting. In summary, after conducting all experiments with various architectures (EfficeientNetV2, VGG16 combined with AutoGluon, BCNN, and DenseNet-121), based on our findings, we implemented the EfficientNetV2 in the publicly available version of our OmicsFootPrint pipeline.

### Feature mapping on the circular image through the polar coordinate system

We developed a polar coordinate system to map omics data features onto a circular image. This system establishes a direct link between the pixels on the transformed circular image and their respective omics features. The reference origin of the polar coordinate system (*O)* was established at the geometric center of the circular image. The reference polar axis was set to coincide with the positive x-axis, which connected *O* to the start of chromosome 1. The reference polar axis served as the baseline of the angle measurement. The angle *θ* was then defined as the angular displacement from the reference polar axis in a clockwise direction. For any given feature *i* (*i*=1, 2,…, M), located on chromosome *N*_*i*_ (*N*_*i*_∈ {1, 2,…, 22, *X*, and *Y*}) at position *pos*i, we calculated the polar angle *θ*_*i*_ as follows:

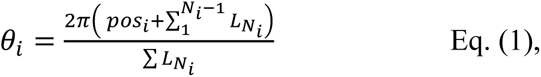

where *L* represents the length of the chromosome *N*_*i*_. This process resulted in a specific feature-angle mapping chart for each type of omics data set, which remained consistent regardless of which specific circular track the omics data set was placed on.

### SHAP Model-Feature Decoding

Here, we used the SHAP values to interpret the trained DNN model by extracting the regions in the circular images (representing omics data) that contributed the most to the classification. By mapping these important regions back to the genome, we identified the specific genomic features and omics data types that were most influential to the model’s classifications.

We calculated SHAP values for every DNN model we trained. As shown in stage 3 of **Figure 1**, these values are presented in a 256×256 matrix for each circular image we input into the model. These SHAP values (log-odds) measure the influence of specific regions on the classification outcome for a given circular image (32). A SHAP value of zero signifies a neutral effect on classification, while positive values indicate a positive contribution to the classification, and negative values suggest the opposite. To identify the most influential regions, we focused on extreme SHAP values that were either above the 95^th^ percentile or below the 5^th^ percentile. We then associated the pixel positions of these significant SHAP values back to the corresponding genomic features by mapping their polar coordinates in the circular images to the previously established feature-angle mapping chart. Specifically, for a given pixel position at column *x*_*i*_ and row *y*_*i*_ in the SHAP output image (*x*_*i*_, *y*_*i*_ = 1, 2,…, 256), we calculated the polar coordinate angle *θ*_*i*_ as:

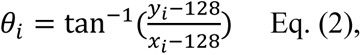

noting that the origin point *O* was set at column 128 and row 128 in the 256×256 SHAP output matrix. By comparing *θ*_*i*_ with the feature-angle mapping chart from Eq. (1), we identified the corresponding omics features. To determine the omics data type, we calculated the polar coordinate radius *R*_*i*_ as:

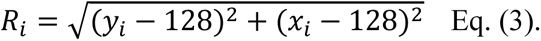

The distribution of radii of all selected pixel positions across all samples was summarized, where peaks were clearly separated by tracks (**Supplementary Figure S1**). The data type of a pixel was then determined by calculating the radius in the polar coordinate system and matching it to the distribution curve peaks.

Theoretically, a feature should be able to be mapped back to the original features accurately. However, the current resolution in the OmicsFootPrint framework restricts the identification of regions of interest to cytobands (resolution 850 bands). Therefore, while the SHAP module identifies broader areas of interest, precise genomic features are inferred based on genomic annotations.

### TCGA multi-omics data collection

In this study, we used lung squamous cell carcinoma (LUSC), adenocarcinoma (LUAD), and breast cancer (BRCA) omics data sets from The Cancer Genome Atlas (TCGA, https://www.cancer.gov/tcga), downloaded through the UCSC Xena datahub (which includes GDC TCGA LUSC, LUAD, and BRCA cohorts) (33). For lung and breast cancer projects, we downloaded copy number data (segmentation format), gene expression data, and RPPA data. For the breast cancer project, we also downloaded microRNA data. The gene expression data included 60,483 gene features as normalized FPKM values. To enhance the signal-to-noise ratio, we focused on the 5,000 most variant genes across the samples in the data set. Breast cancer PAM50 subtype information was obtained from the R package TCGAbiolinks (34). Histologic subtypes of the TCGA BRCA cohort were obtained from a study by Ciriello et al. (35).

### Benchmarking Implementation

To evaluate the performance of OmicsFootPrint, we benchmarked the model’s AUC with publicly available multi-omics data sets containing seven train/test combinations for six different drugs across three drug response databases (GDSC, TCGA, and PDX) (13). The data sets contained three types of omics data: gene expression, CNV, and somatic mutation. The detailed sample sizes for each of the tested data sets are provided in **Supplementary Table 5**. For benchmarking, we evaluated two advanced deep-learning algorithms. The first is DeepInsight-3D, an approach that utilizes image-based data. The second is MOLI, which employs a method based on analyzing tabular data. The best performance AUCs for these deep-learning algorithms with the same data set were obtained from a recent publication (22). In addition, we evaluated two unsupervised machine learning algorithms based on the feature selection strategies, non-negative matrix factorization (NMF) and integrative NMF (intNMF). For both methods, features were scored following the methodology of Kim et al. (36), such that those features exceeding the median plus three mean absolute deviations were retained for the factorization of the independent data sets. The AUC of the clustering performance (k=2) was evaluated against the observed drug response.

We also benchmarked OmicsFootPrint against five supervised machine learning algorithms (kernel SVM, NNLS, LDA, Ranger, and GLMnet), training the models with 10-fold cross-validation as implemented in R package SuperLearner. To provide a comprehensive evaluation of OmicsFootPrint’s performance, we compared the range of the machine learning algorithm’s ability to predict independent data sets. All algorithms were assessed with default settings (13). The results for DeepInsight-3D and MOLI were obtained directly from the study by Sharma et al. (22).

### Bioinformatics and Statistics Analysis

Enrichment analyses of the features identified in this study were conducted using Fisher’s exact test. This test was applied to feature frequency counts. We assessed the predictive performance of various models constructed by permuting the datasets through the analysis of area under the curve values. To accommodate the repeated partitions applied across comparisons, we employed repeated measures ANOVA. Specifically, a one-way repeated measures ANOVA was utilized for the analysis of the two-class TCGA lung cohort, while a two-way repeated measures ANOVA was applied for the four-class TCGA breast cohort. Following the ANOVA, post-hoc paired t-tests were conducted to examine the main single effect, allowing for a detailed comparison of model performances. Differential expression analyses were performed with the R package limma. Annotation-based pathway enrichment analyses were performed using the R package gprofiler2 (37). Network construction was carried out through QIAGEN Ingenuity Pathway Analysis (38).

### Software Implementation and Code Availability

The generation of circular images in the study was performed using the circlize package in R (39). All deep learning models were implemented in Python (version 3.9) with TensorFlow framework (version 2.10.0). Additional Python packages were utilized to support different aspects of the analysis, including scikit-learn (version 1.2.2), pandas (version 1.5.3), matplotlib (version 3.5.0), and Pillow (version 9.4.0). The SHAP Python package/library was used to implement the determination of SHAP values for a specifically trained DNN model (40). All codes and scripts used in this study are available at https://github.com/KalariRKLab-Mayo/OmicsFootPrint.

## RESULTS

We developed the novel deep-learning framework OmicsFootPrint to convert patient-level multi-omics data into 2-D circular images for each patient for image classification (**Figure 1**). An individual omics data type is represented in a track in circular layouts, with the arrangement of data types with a higher number of features placed on the outer tracks. In contrast, those with fewer features were located on the inner tracks. **Figure 2A** shows an image representation of the omics data, including gene expression, copy number variations, and data types in individual tracks. A transformed circular image where data points along the circumference represent singular omics features is used as the input into the OmicsFootPrint framework for subsequent classification (**Figure 2B)**. An example of the 700 individual samples OmicsFootPrints arranged in a grid, highlighting the capability of our framework to discover patterns and facilitate robust multi-sample analysis, is shown in **Figure 2C**.

The circular image sets were input into the DNN model (**Figure 1**), which leveraged transfer learning techniques using the pre-trained weights from the ImageNet for initialization. Subsequent SHAP decoding was incorporated into the OmicsFootPrint framework to indicate how each feature contributes to the model’s output, as detailed in the Methods section. Key identified features were then aggregated across samples to gain a deeper phenotypic understanding of the cohort (**Figure 1**, stage 3).

### Evaluating the OmicsFootPrint framework with large patient data sets

We utilized the lung and breast cancer data cohorts from TCGA to conduct real-world evaluations of the OmicsFootPrint framework. For each classification test, we selected sample cohorts with complete data (copy number, gene expression, RPPA, and/or microRNA). (1) First, we evaluated the performance of the OmicsFootPrint framework using four different architectures, focusing on circular images derived from gene expression data only. (2) Second, we assessed the performance of the OmicsFootPrint with the best-performing architecture across single, double, and multi-omics data types in two-class and multi-subtype classification contexts. For the two-class classification, we utilized 682 TCGA lung cancer samples (359 LUAD and 323 LUSC). For the multi-subtype classification, we utilized 836 TCGA BRCA molecular subtypes (PAM50 – a gene signature derived using 50 genes), composed of 434 Luminal A, 175 Luminal B, 153 Basal, and 74 HER2 samples. Normal-like PAM50 calls from the TCGA BRCA cohort were excluded as they represent potentially low-quality tumor samples. (3) Finally, we conducted a feature selection analysis using SHAP values to investigate the histological differences of 493 TCGA BRCA samples, consisting of 400 breast-invasive ductal carcinoma samples (IDC) and 93 breast-invasive lobular carcinoma (ILC) samples. All the data sets used in this study are summarized in **Table 1**.

**Table 1:** TCGA data sets and sizes evaluated with the OmicsFootPrint framework. Abbreviations: BRCA, breast cancer; LUAD, lung adenocarcinoma; LUSC, lung squamous cell carcinoma; TCGA, The Cancer Genome Atlas.

| Data Cohort and Subtypes |  | Cohort Sizes |
| --- | --- | --- |
| TCGA Lung | **Total** | 682 |
|  | LUAD | 359 |
|  | LUSC | 323 |
| TCGA BRCA PAM50 | **Total** | 836 |
|  | Luminal A | 434 |
|  | Luminal B | 175 |
|  | HER2 | 74 |
|  | Basal-like | 153 |
| TCGA BRCA histologic | **Total** | 493 |
|  | Ductal | 400 |
|  | Lobular | 93 |

### EfficientNetV2 was the best-performing DNN architecture in OmicsFootPrint classification

The OmicsFootPrint framework can utilize various deep-learning models as its backend. We evaluated the classification performance of OmicsFootPrint with several state-of-the-art DNN architectures, including BCNN, DenseNet-121, EfficientNetV2, and AutoGluon with a feature vector (VGG16+AutoGluon). To simplify, we assessed with single-omics circular plots using gene expression data. Ten random partitions were generated for the TCGA lung cohort (LUAD and LUSC classification) and TCGA breast cancer data sets (consisting of four molecular subtypes defined using the PAM50 method), respectively.

Figure 3A illustrates the performance of the four DNN architectures in the TCGA lung cohort. All four architectures showed high performance (median AUCs ≥ 0.90 and minimum AUC=0.84). Among them, three architectures (EfficientNetV2, DenseNet121, and Bilinear) achieved high AUCs (median AUC > 0.95). EfficientNetV2 outperformed DenseNet121 and Bilinear, with the highest median AUC of 0.98, which was not statistically significant (**Table 2** and Supplementary Table S1). In the more complex four-subtype BRCA data cohort, we observed that the performance of DNN architectures varied with BRCA subtypes (Figure 3B). In the basal-like subtype, all four DNN architectures showed good performance (all AUCs > 0.8 except one model from VGG16+Autogulon). It was the most challenging in the Luminal B subtype, where only one model from EfficientNetV2 achieved an AUC above 0.8. Nevertheless, EfficientNetV2 showed the highest number of trained models with good performance (AUCs>0.8) out of the 10 repetitions for all four subtypes. Paired t-test showed that EfficientNetV2 outperformed the other three architectures across all BRCA subtypes significantly (adjusted p-values: 0.01 vs. Densenet-121, 0.006 vs. Bilinear, and 7.3x 10-6 vs. VGG16+Autogulon) (**Table 2** and and Supplementary Table S2).

**Figure 3.**
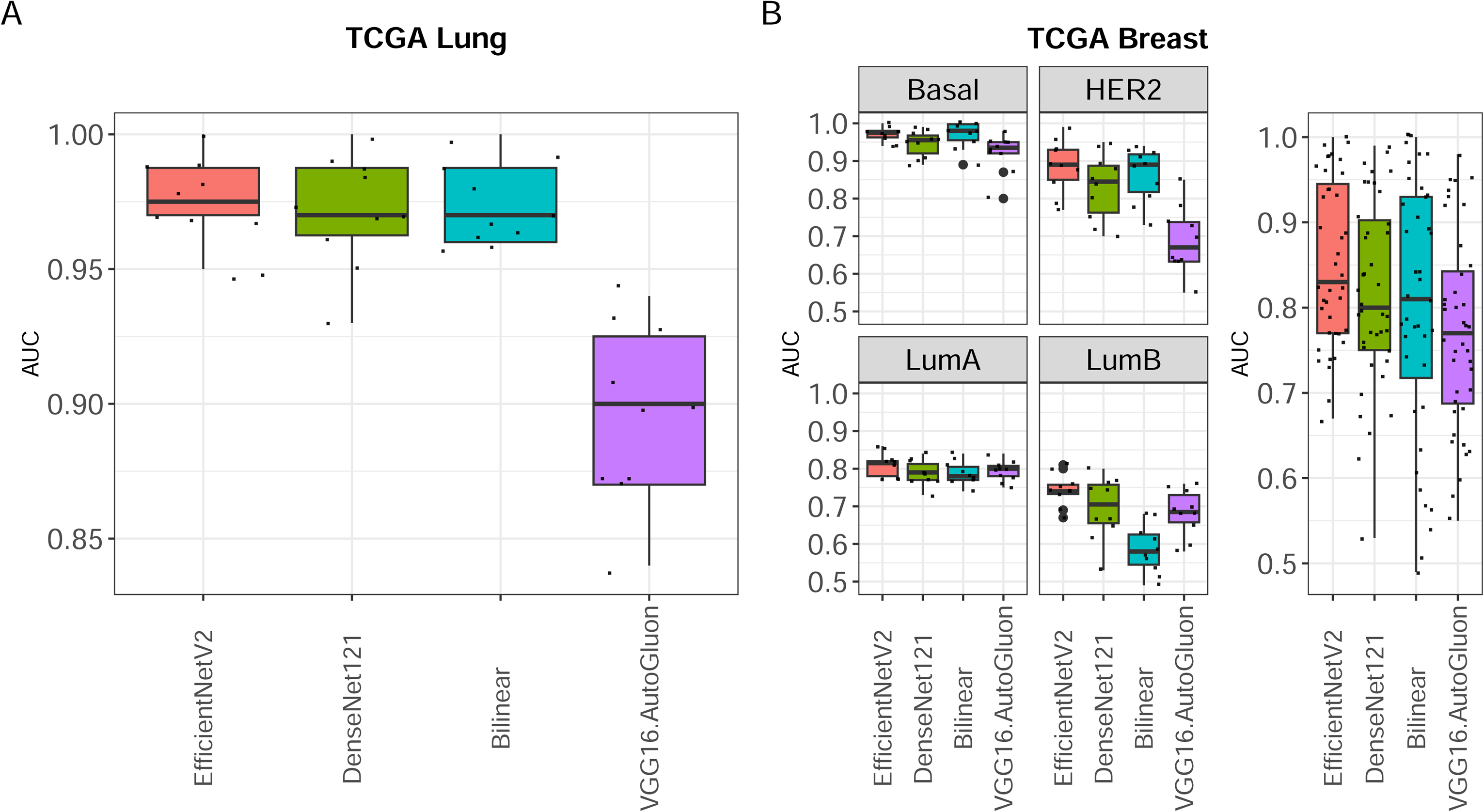
DNN architecture comparison. We evaluated different DNN architectures in TCGA Lung (A) and BRCA PAM50 (B) data cohorts. EfficientNetV2 showed the best performance in both cohorts.

**Table 2:** Performance metrics of the 4 neural network implementations with the OmicsFootPrint framework (numbers in the table showed median AUC± standard deviation).

|  | Lung | Breast |  |  |  |  |
| --- | --- | --- | --- | --- | --- | --- |
|  |  | Luminal A | Luminal B | Basal | HER2 | Overall |
| **EfficientNet_V2** | **0.98±0.02** | **0.82±0.03** | **0.74±0.04** | **0.98±0.02** | **0.89±0.07** | **0.83±0.07** |
| **DenseNet-121** | 0.97±0.02 | 0.79±0.03 | 0.71±0.08 | 0.96±0.03 | 0.85±0.09 | 0.80±0.11 |
| **Bilinear** | 0.97±0.02 | 0.78±0.03 | 0.58±0.07 | 0.98±0.04 | 0.89±0.07 | 0.81±0.15 |
| **VGG16+Autogluon** | 0.90±0.03 | 0.80±0.03 | 0.69±0.06 | 0.94±0.05 | 0.67±0.09 | 0.77±0.11 |

### Multi-omics data showed the best prediction power in TCGA Lung and Breast cancer subtype classification

To assess the predictive capability of various combinations of omics data, we conducted a performance evaluation of OmicsFootPrint sets created using three different data combinations: single-omics (gene expression only), two-omics (gene expression + CNV), and multi-omics (gene expression + CNV + protein and phosphoprotein data from RPPA assay). We generated image sets for all three data combinations by employing ten random splits and subsequently processed them through OmicsFootPrint, using EfficientNetV2 as the backend DNN model.

Figure 4A summarizes the resulting AUC values for each data combination in the context of TCGA lung cancer. We observed that the multi-omics data (gene expression, CNV, and RPPA) showed higher performance when compared to the other data combinations (expression only or expression + CNV data) (**Supplementary Table S3**). In general, in the lung cancer cohort, we noted that all data combinations yielded high AUC values (median AUC=0.92 and minimum AUC=0.87). Expression + CNV + RPPA showed the highest median AUC of 0.94, while 0.92 for expression + CNV and 0.91 for expression only. No statistically significant difference was detected among the data combinations (P=0.24 by repeated measures ANOVA). This observation suggests that gene expression data alone may suffice for distinguishing between LUSC and LUAD within this lung TCGA cohort.

We subsequently investigated a more challenging task involving multi-class classification using the TCGA BRCA cohort, which contains four PAM50 molecular subtypes. This classification task was performed using four different data combinations: single-omics (expression only and CNV only), double-omics (expression + CNV), and multi-omics (expression + CNV + protein and phosphoprotein from RPPA). We omitted RPPA in the single and double omics data combinations due to its sparsity, which was caused by a limited number of protein and phosphoprotein measurements. In contrast to the two-subtype lung cancer cohort classification, we observed greater variability in performance in the four-subtype classification task (Figure 4B). We observed a similar trend in architecture selection, where higher performances were observed in basal-like subtypes and lower in luminal subtypes. Among the omics combinations, the multi-omics image sets (expression + CNV + RPPA) demonstrate the most robust top performance across different subtypes (highest median AUC shown in bold fonts in **Supplementary Table S4**). The model trained with multi-omics image sets showed the highest overall median AUC of 0.86, although not as significant as expression only and expression + CNV. In summary of all comparisons for both lung and breast cohorts, we chose the multi-omics image sets that demonstrated the best model performance.

**Figure 4.**
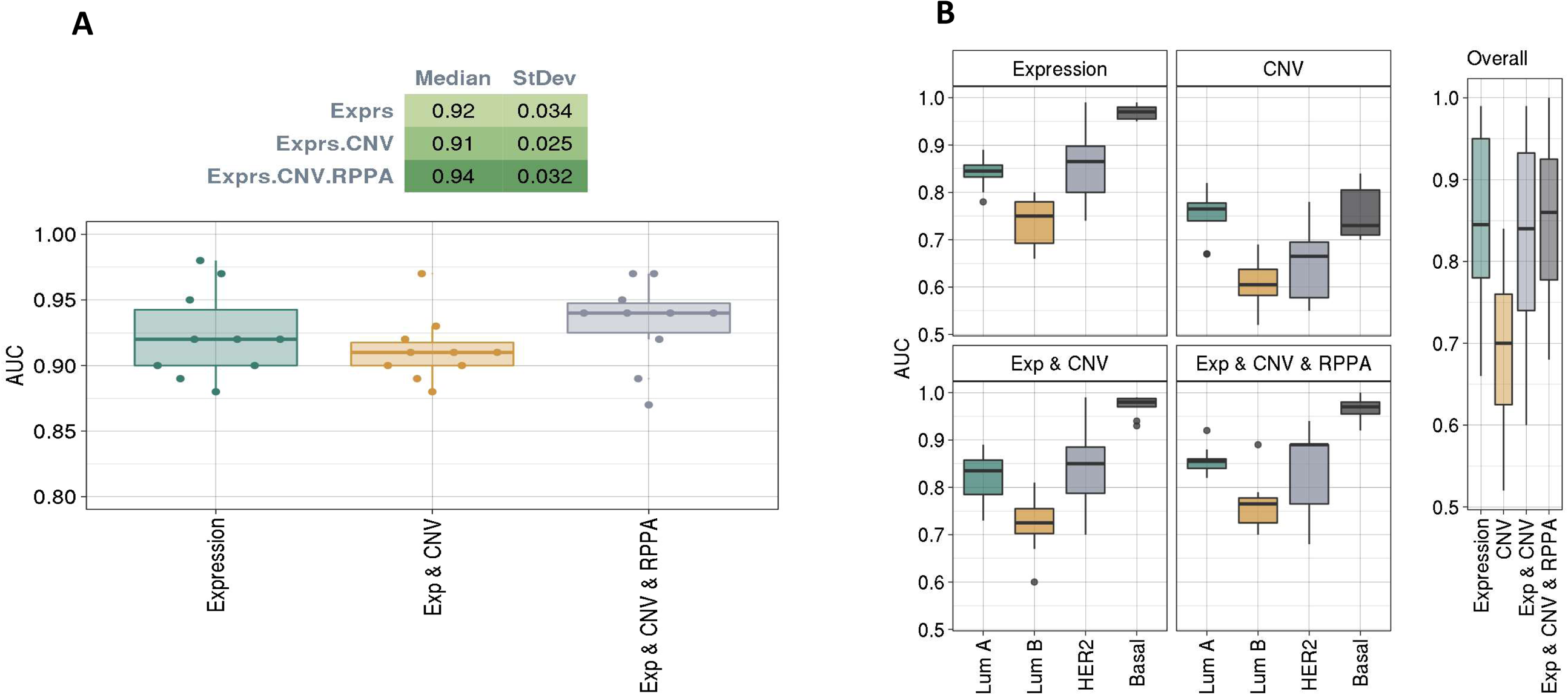
Comparison of different omics combinations. We evaluated different combinations of omics data in TCGA Lung (A) and BRCA PAM50 (B) data cohorts. The integration of multiomics data showed the best performance in both cohorts, while expression data alone showed good performance.

### Benchmarking OmicsFootPrint against top deep learning and machine learning algorithms in drug response prediction using cancer cell lines data

To evaluate the performance of the OmicsFootPrint framework, we conducted a benchmarking comparison with nine methods against a set of public multiomics datasets of three omics datatype (gene expression, CNA, and somatic mutation) with drug response data (TCGA, GDSC, and PDX). DeepInsight-3D is a deep learning image-based approach for analyzing multi-omics data, and MOLI is a deep learning method using tabular data. Additionally, traditional machine learning algorithms, including NMF, intNMF, SVM, Ranger, GLMnet, NNLS, and LDA (See Methods), were evaluated.

The AUC scores for all drug response data sets and algorithms are summarized in Figure 5. The overall performance of all benchmarking tests (across drugs and different algorithms) showed a relatively low average AUC of 0.57, which is primarily attributable to the complexity of drug response prediction and the presence of a highly imbalanced training and testing data set (**Supplementary Table S5**) in the cancer cell lines database. All three deep learning algorithms, such as MOLI, DeepInsight-3D, and OmicsFootPrint, outperformed (average AUC of 0.67) the conventional machine learning algorithms with an average AUC of 0.52. Among the three deep learning algorithms, OmicsFootPrint performed best in 5 out of 7 drug response studies and achieved the highest average AUC across all data sets (0.74).

**Figure 5.**
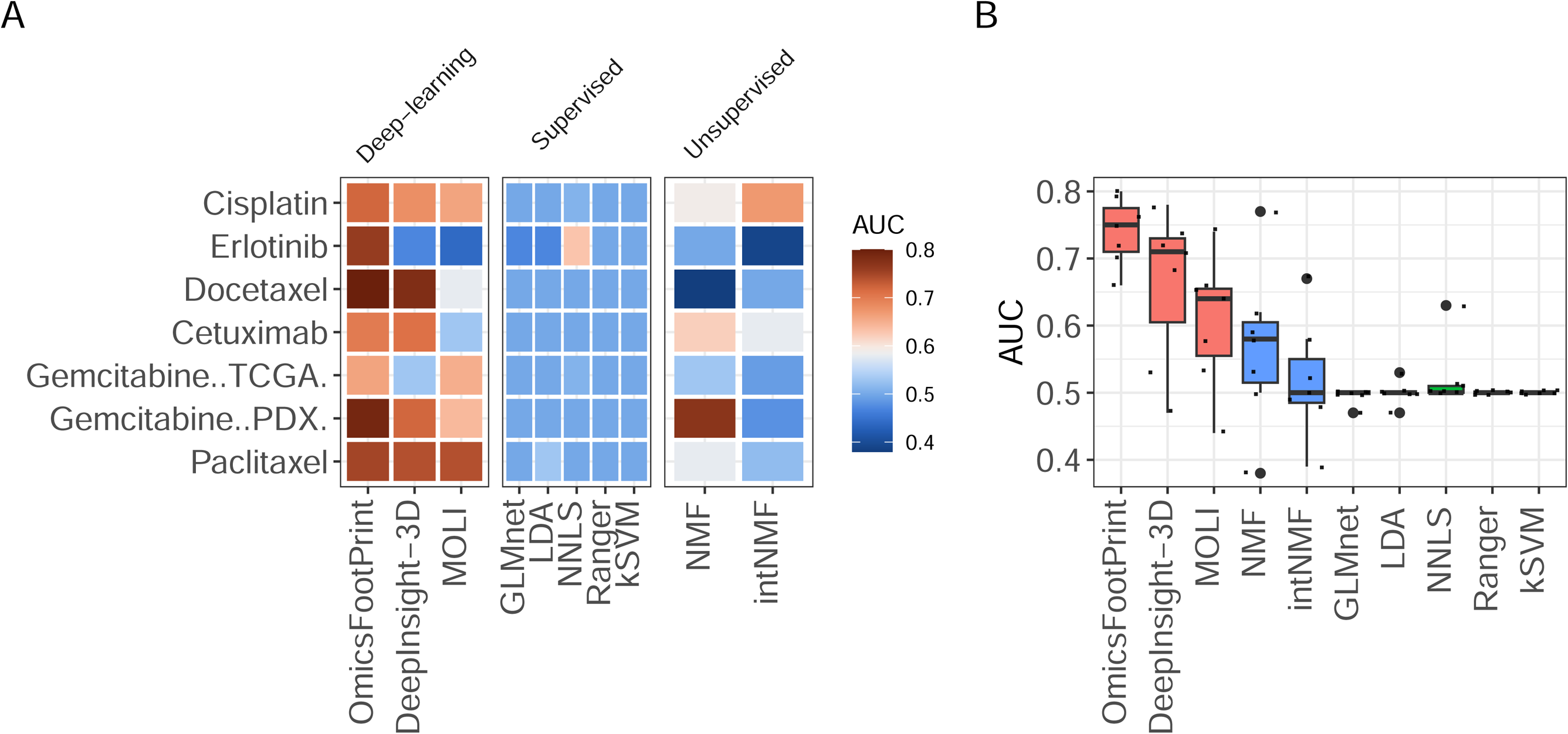
Benchmarking of OmicsFootPrint with other state-of-the-art multiomics integration methods in predicting drug response in a public data set. Metrics were evaluated with AUCs. OmicsFootPrint showed the best performance in 5 out of 7 drug-response data sets.

### Comprehensive multi-omics analysis and histologic subtype classification in breast cancer using the OmicsFootPrint framework

The OmicsFootPrint framework is not just a tool for classification using deep learning; it also interprets which multi-omics features are most influential in making predictions for a given sample. It does this by identifying key coordinates in circular images using SHAP values and then linking these coordinates back to their respective omics features. This approach allows for the analysis of multi-omics networks by examining the co-occurrence of various omics features. We demonstrate this with an example where we classify and compare different histological subtypes of breast cancer using data from the TCGA cohort.

We applied the OmicsFootPrint method on IDC and ILC, two major histological subtypes of breast cancer, using the TCGA BRCA data set. We conducted our analysis using both a three-omics and a four-omics data design. In the three-omics approach, we utilized gene expression, CNV, and RPPA data across 493 samples. For the four-omics approach, we extended the analysis by incorporating miRNA data; this reduced our sample size slightly to 480 samples. The performance of the predictive models was similar between the three-and four-omics designs, with average AUC values of 0.87±0.06 and 0.87±0.04, respectively.

Next, we demonstrated the performance of the SHAP decoding module using the four-omics data set. For each sample, SHAP values were generated based on the trained model. We focused on the most significant SHAP values, identifying positive peaks (values above the 95th percentile) and negative peaks (values below the 5th percentile). These peak values were then correlated back to corresponding omics features using their polar coordinates, as detailed in the Methods section. A total of 251 omics features were identified as significantly enriched. Of these, 145 features (57.8%) were identified by only positive peaks, 71 features (28.3%) were identified by only negative peaks, and 35 features (13.9%) were identified by both positive and negative peaks ("double-peak" features). These double-peak features were observed across all four omics data types (**Table 3**), including many reported to be associated with ILC (Figures 6A **and 6B**). For example, gene expression and CNV features at 16q22.1 harbor the *CDH1* gene, which is known to be crucial in differentiating in ILC due to its copy number loss and lower expression in comparison to IDC. Additionally, RPPA features at 10q23, harboring probes for *PTEN*, were reported to show lower protein expression in ILC than in IDC. Our functional enrichment analysis of all double-peak features against the REACTOME pathway database showed significant enrichment of *PI3K/AKT* signaling in cancer (adjusted p-value=0.0072). Besides these known genes, we identified miRNA features at 22q13 covering 19 significant miRNA probes, with hsa-mir-33a being most differentially expressed (adjusted p-value=3.9×10^-15^). MiR33a is known to target *ZEB1*, a regulator of *CDH1* and cell adhesion (41), suggesting a mechanism of dysregulated cell-cell adhesion in ILC (Figure 6C).

**Figure 6.**
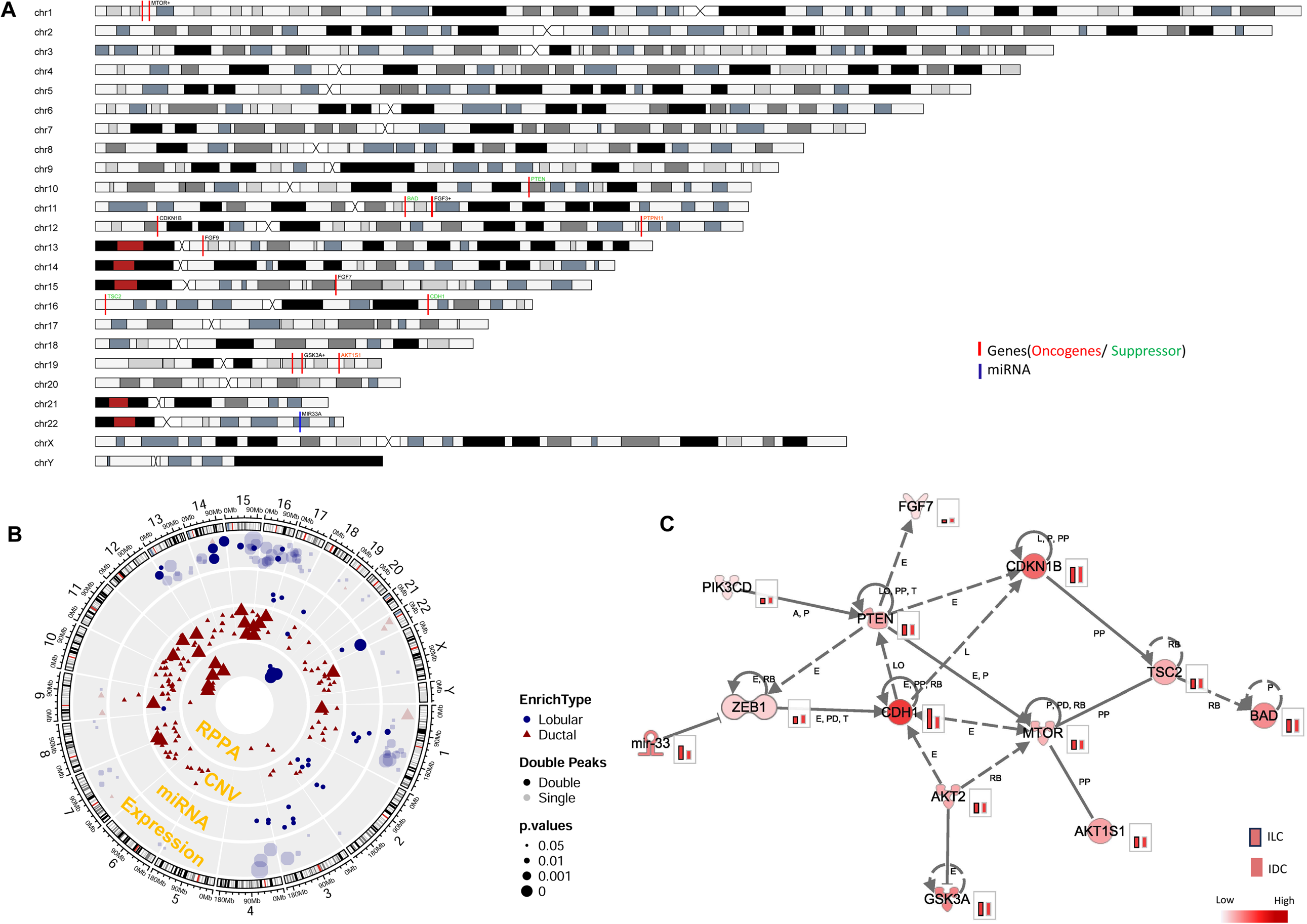
In the analysis of four omics datasets from the TCGA BRCA cohort, the OmicsFootPrint framework successfully distinguished between invasive lobular carcinoma and invasive ductal carcinoma, highlighting key genes such as CDH1, PTEN, and AKT. The results are presented in several formats: (A) Gene features identified by OmicsFootPrint are shown in an ideogram, illustrating the genomic landscape. (B) These features are further visualized in a circular image, where different shapes and colors denote enriched subtypes; larger symbols indicate features with double-peak patterns, and varying color opacity reflect the significance of the p-values. (C) Further examination of the gene features suggested that miR33a might influence the histological differences between the two subtypes, showing potential novel mechanisms (the color scale representing gene expression levels for ILC (framed) and IDC (unframed)).

**Table 3:**
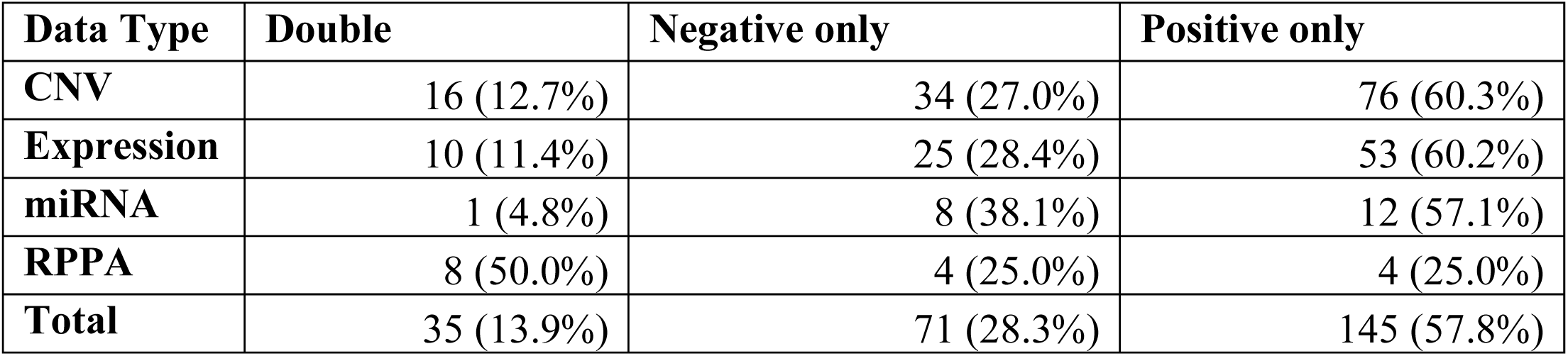
Summary of significant differentiated features Abbreviations: CNV, copy number variation; miRNA, microRNA; RPPA, Reverse Phase Protein Array.

| Data Type | Double | Negative only | Positive only |
| --- | --- | --- | --- |
| CNV | 16 (12.7%) | 34 (27.0%) | 76 (60.3%) |
| Expression | 10 (11.4%) | 25 (28.4%) | 53 (60.2%) |
| miRNA | 1 (4.8%) | 8 (38.1%) | 12 (57.1%) |
| RPPA | 8 (50.0%) | 4 (25.0%) | 4 (25.0%) |
| Total | 35 (13.9%) | 71 (28.3%) | 145 (57.8%) |

### OmicsFootPrint model remained robust with sample size variation and track arrangement

OmicsFootPrint exhibits the capability to operate effectively with relatively small sample sizes through the application of transfer learning from the pre-trained ImageNet database. This feature is particularly advantageous when dealing with high-dimensional, low-sample-size multiomics clinical data sets from real-world studies. To assess the impact of training set size on OmicsFootPrint’s classification performance, we conducted a simulation test by gradually reducing the sample size. We tested on the TCGA lung cancer cohort because it consists of balanced sample sizes across the two subtypes. We systematically decreased the size of the training multi-omics image set from 100% (N=470, LUAD: LUSC=253:217) to 10% (N=46, LUAD: LUSC=25:21) in 10% decrements each with 10 permutations. The resulting AUC remained ≥ 0.95 for all tests until the training set was reduced to less than 20% (LUAD: LUSC= 51:43) (Figure 7). When the training set was reduced to 10%, OmicsFootPrint still produced an AUC of 0.86; however, there also was a notable increase in performance variability (ranging from 0.77∼0.92). This observation highlights OmicsFootPrint’s resilience when working with a small training data set.

**Figures 7.**
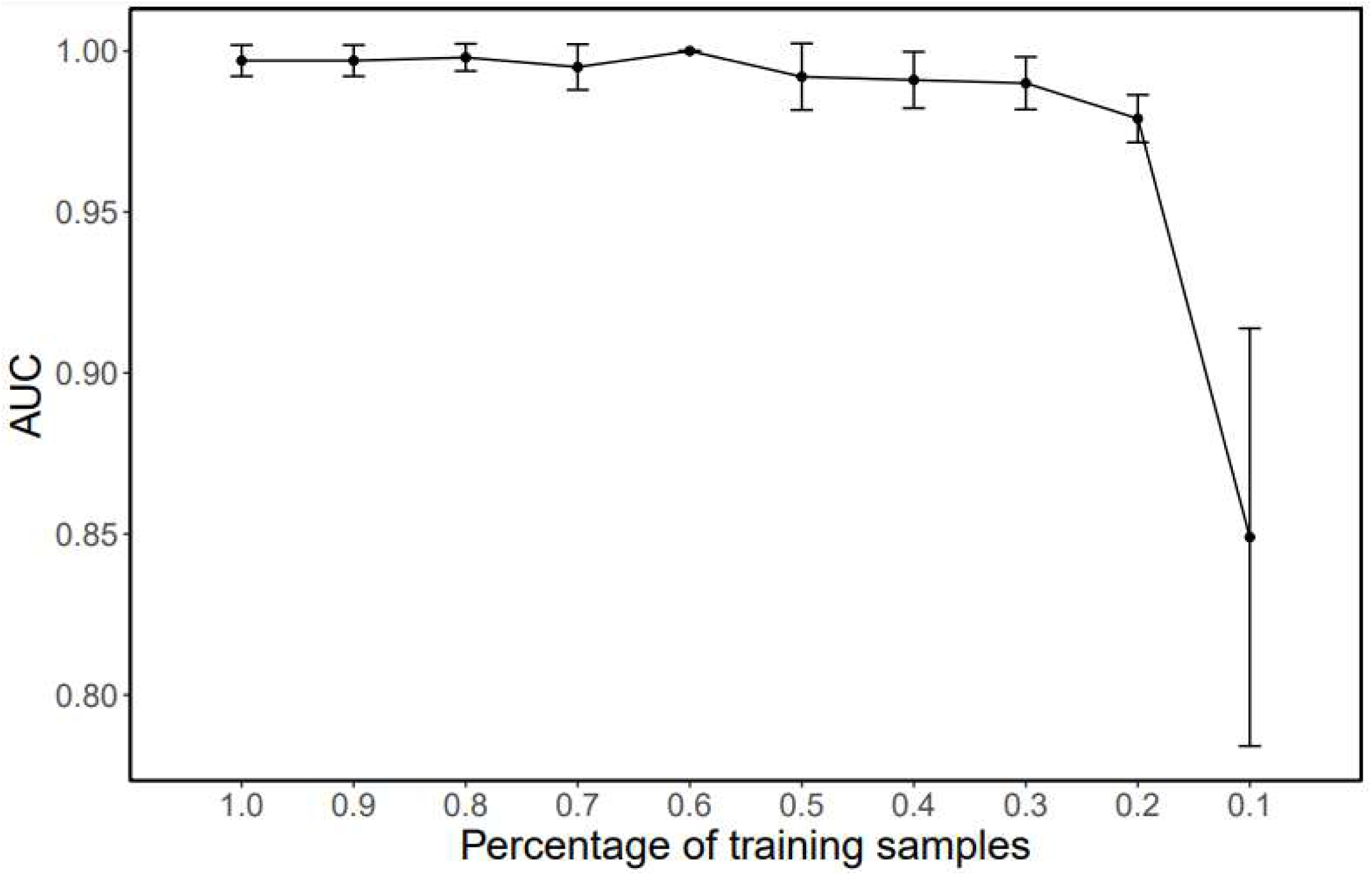
Robustness of the OmicsFootPrint framework with reduced training sample sizes. The figure displays the performance of the OmicsFootPrint in relation to varying training sample sizes within the TCGA Lung data cohort. The y-axis represents the area under the curve as a measure of model performance, while the x-axis indicates the percentage of the total available data used for training. The AUC values remain consistently above 0.95, showing the model’s high predictive accuracy, until the training sample size decreases to below 20%. This demonstrates OmicsFootPrint’s robustness against reductions in training data quantity.

We also examined the arrangement of omics tracks within the same TCGA lung cancer multi-omics data set. Three different arrangements listed from the outermost to innermost track were tested: Expression/CNV/RPPA, CNV/Expression/RPPA, and CNV/RPPA/Expression listed from the outermost to innermost track. Our analysis, consisting of 10 repetitions, did not show any significant differences in performance among the various track arrangements (P=0.66 by ANOVA). OmicsFootPrint consistently showed stable classification performance regardless of the track arrangements.

### Computational Resource and Performance

In the current setup of the OmicsFootPrint, each circular image is created with a resolution of 1024×1024 pixels in PNG format, and the file size for each image ranges from 20 KB to 150 KB. For this study, our primary computing resource was a high-performance computing node equipped with two NVIDIA V100 GPUs, each with 32 GB of memory. On average, training sessions consisted of 100 epochs with a batch size of 32, and each session took around 3 hours to complete.

## DISCUSSION

In this study, we introduced a new framework, OmicsFootPrint, which employs an image-based deep learning model for integrated multi-omics analysis (Figure 1). The core concept involves transforming multi-omics data into two-dimensional circular images for each sample. In these images, the outermost layer consistently represents the chromosome, providing a fixed reference point. Inside layers represent omics data, plotted on separate tracks within the circular plot (Figure 2). The gene annotations and data specific to each omics platform guide the placement of data within these inner tracks. Our approach allows for image classification using advanced deep-learning tools. Model results are explained by identifying informative features using the SHAP method.

Different omics data types can be incorporated with this circular arrangement. Continuous omics data types, including gene or miRNA expression from RNA-seq or microarray, CNV from DNA sequencing, protein abundance measured through RPPA assays or mass spectrometry, and epigenomics data, including methylation, CHIP-seq, ATAC-seq, and others, can be included. Normalized data are suggested here, although the fixed circular track heights for all samples perform intrinsic standardization. While this study did not include categorical omics data like point mutations, it is thought that they could be easily integrated into this methodology by translating them into discrete numeric states or ratios of the alternate allele to reference allele. Such an approach would allow for comprehensive and integrative analysis that includes both continuous and categorical omics data.

We first compared different compositions (single-omics, double-omics, and multi-omics) of the circular images in the classification of LUSC and LUAD subtypes in the TCGA lung cancer cohort. It is observed that gene expression data were sufficient for this comparison, which is not surprising as transcriptomic data are widely acknowledged as an information-rich data set. The multi-omics model incorporating expression with copy number and RPPA presented a slight increase in performance (Figure 3).

The OmicsFootPrint framework has been developed with adaptability in mind, ensuring compatibility with a wide range of DNN architectures. We compared the performance of several state-of-the-art DNN models (VGG+AutoGluon, BilinearCNN, DenseNet-121, and EfficientNetV2) in the classification of PAM50 subtypes in the TCGA breast cancer cohort (Figure 3). In this case, the OmicsFootPrint was not limited to binary classification; instead, it was trained and utilized to learn and distinguish between the four different subtypes concurrently. As shown in Figure 3, the EfficientNetV2 performed best overall and had the most consistent performance in AUC across the lung and breast cancer datasets and subtypes. In breast cancer, we observed that the basal-like samples presented the best discriminatory power, with all four DNN models showing AUC above 0.90. The HER2 breast cancer subtype, while presenting as the most under-represented (n=74), was also well discriminated (AUC > 0.85) except for VGG+AutoGluon. Luminal B was the most challenging subtype for identification, likely due to the high similarity to the luminal A subtype, which are both hormone receptor (HR) positive derived from luminal epithelial cells (42,43). In contrast, the HER2 cancers are molecularly distinct and driven by a 17q12 amplification containing the ERBB2 gene, and the basal-like cancers have been proposed to originate from basal or myoepithelial cells (44).

We then conducted a benchmarking data analysis using the OmicsFootPrint framework against a range of multi-omics drug-response data sets from cancer cell lines. This comparison included other deep-learning algorithms such as DeepInsight-3D and MOLI, unsupervised learning algorithms like NMF and intNMF, and several standard machine-learning algorithms, including SVM, Ranger, GLMnet, NNLS, and LDA. In these tests, the deep learning algorithms, including OmicsFootPrint, generally outperformed the machine learning algorithms. OmicsFootPrint achieved the best results in 5 out of the 7 drug-response data sets (Figure 5). In two particularly imbalanced data sets, it was slightly outperformed by DeepInsight-3D, placing second. These outcomes highlight OmicsFootPrints significant potential in tackling complex studies, such as drug-response prediction. A key distinction between OmicsFootPrint and DeepInsight-3D lies in their capacity to analyze multi-omics data. DeepInsight-3D is limited to handling three omics data sets, whereas OmicsFootPrint can manage a broader range. We illustrated OmicsFootPrint’s capabilities with a comparative analysis of invasive lobular and ductal carcinomas in the TCGA BRCA cohort. By employing four distinct omics data types, OmicsFootPrint effectively demonstrated its robust performance in accurately classifying these histologic subtypes (Figure 6). This application was focused on samples with available histologic clinical information, encompassing a total of 493 cases. The average AUC of this subtype classification was 0.87±0.06. In general, the challenges in classification accuracy might be attributed to several factors: the unbalanced sample sizes of IDC and ILC (400:93), the complexity arising from clinical subtype heterogeneity, and the high degree of molecular similarity between the subtypes.

The OmicsFootPrint framework consists of a novel approach to interpreting the predictions of multi-omics data from a trained neural network using SHAP values. These values assign an importance score to each image prediction, allowing us to map the top SHAP values back to their corresponding omics features. This effectively identifies the most critical features distinguishing between subtypes/groups. We demonstrated this with an explainable multi-omics integration using four types of omics data (gene expression, copy number, miRNA, and RPPA) to evaluate histological differences (IDC vs. ILC) in breast cancer. Our analysis focused on highly predictive samples (prediction probability > 0.95 with concordant classification), identifying 251 features significantly enriched in one of the histological subtypes. We paid particular attention to ‘double-peak’ features, which showed significance in both positive and negative SHAP peaks. OmicsFootPrint successfully pinpointed key molecular differences between ILC and IDC. For instance, *CDH1* copy number and gene expression at 16q22 were notably significant in differentiating the two subtypes, aligning with *CDH1*, a well-known marker of ILC (45). Additionally, *PTEN* RPPA at 10q23 was identified, and it is known for its lower protein expression in ILC compared to IDC. The *PI3K/AKT* signaling pathway, significantly enriched in the double-peak features, was reported to be more active in ILC than in IDC. Furthermore, among the significant miRNA features at 22q13, miRNA-33a showed a marked differential expression between the subtypes (Figure 6). Previous studies have linked MiR33a to the regulation of *CDH1* and cell adhesion through *ZEB1*, suggesting a potential mechanism for disrupted cell-cell adhesion in ILC (41). In summary, the capability of OmicsFootPrint to simultaneously analyze four different omics data types has showcased its superior performance compared to existing methods like DeepInsight-3D. Its innovative approach to image-based integration and analysis of multiomics data offers a powerful tool for revealing complex biological insights.

A key advantage of OmicsFootPrint as a deep learning method is its use of transfer learning. It adopts pre-trained weights from ImageNet, an extensive visual database containing over 14 million images (24). This strategy enables OmicsFootPrint to handle smaller sample sizes effectively, a significant deviation from the traditional deep learning approaches that often require large amounts of samples for decent performance in multi-omics integration analyses without being overfitting (18). Our simulations on sample size reduction demonstrated that OmicsFootPrint could maintain robust performance (AUC > 0.95) with fewer than hundreds of images per subtype group. This level of efficiency makes it highly practical for many clinical studies in multi-omics settings.

Another advantage of OmicsFootPrint is its ability to convert traditionally large multi-omics files into more compact images, significantly reducing memory requirements. For instance, in the GDC data portal for a single TCGA patient, the combined file size for gene expression, copy number, and RPPA data can amount to approximately 5 MB each. In contrast, a circular image created by OmicsFootPrint, even at a high resolution of 1024×1024 pixels, occupies only about 100 KB. This compact format enables the use of single-sample multi-omics circular images as concise representations of a patient’s multi-omics landscape. These images can be seamlessly integrated into personal electronic medical records (EMR), making them highly practical for personalized precision medicine due to their large information capacity and relatively small storage footprint.

There are limitations to the current design of the OmicsFootPrint framework. First, the current OmicsFootPrint rescales the original circular image to 256×256 pixels to facilitate transfer learning from the ImageNet, which causes inevitable information loss during compression. The reduced resolution also causes difficulties in mapping SHAP values back to the genomic features. The current limitation for gene decoding is limited to the cytoband level, which needs further mapping to the original omics data for gene-level information. The second limitation is the restricted track space, which leads to a down-selection of features for specific omics data to improve the signal-to-noise ratio. For example, whole transcriptome gene expression data (either from microarray or RNA-seq) generally have 20,000∼60,000 gene features based on the probes or annotation used. However, many of these genes are either not expressed or show slight variation across samples. Removing such "background noise" features is crucial to improving the clarity and informational value of the multi-omics images. Despite these limitations, we see significant opportunities for improvements. In the next version of OmicsFootPrint, we plan to revamp the circular images by utilizing RGB colors or grayscale to represent feature values instead of the current black-and-white scheme. This change is expected to significantly increase the density of information conveyed, enabling the inclusion of additional omics data types (e.g., mutations). Moreover, rather than downsizing the images and risking the loss of information, we intend to segment the circular images into 256×256 pixel patches for processing at much higher resolutions. This approach should substantially refine the resolution for SHAP deconvolution, leading to more detailed and informative analyses.

## CONCLUSION

We developed a novel image-based deep learning algorithm, OmicsFootPrint, for integrative multi-omics analysis. We benchmarked with state-of-the-art deep-learning and machine-learning algorithms and showed superior performance of OmicsFootPrint. We also demonstrated how the SHAP-based explanation module of OmicsFootPrint could help interpret the trained DNN model and indicate how much each feature contributes to the model’s output. These features could help elucidate the underlying mechanisms of disease.

## AUTHOR CONTRIBUTIONS

X.T. and N.P.contributed equally to the manuscript.

X.T., N.P., K.J.T., R.W., C.C.O., J.C.B., H.R.T., E.W.K., L.W., M.P.G., V.S. and K.R.K. conceived the idea and reviewed the manuscript. X.T., N.P., K.J.T., V.S., and K.R.K. conducted analysis and prepared the manuscript. All authors contributed to the discussion, and overall data interpretation and approved the final manuscript. All authors read and approved the final manuscript.

## ACKNOWLEDGMENTS

We acknowledge the use of ChatGPT, a conversational artificial intelligence tool developed by OpenAI, to assist in clarifying and refining the sentences in this manuscript based on the material we provided. The Scientific Publications staff at Mayo Clinic provided copyediting support.

## Data Availability Statement

The complete circular image data sets for this study can be found in the zenodo (reference to be determined). Deidentified sample image sets were provided at https://github.com/KalariRKLab-Mayo/OmicsFootPrint.

## Supplementary Information

**Supplementary Figure S1**: Distribution of the distances of peaks to the center of the image. The density curve shows that peaks are enriched by data type. This property is used to determine the omics data type of any peak point on the circular image in a data-driven way.

**Supplementary Table S1:** Comparison of AUCs across different architectures for different subtypes in the lung cohort.

**Supplementary Table S2:** Comparison of AUCs across different omics combination for different subtypes in the breast cancer cohort.

**Supplementary Table S3:** Comparison of AUCs across different omics datatype combinations for different subtypes in the lung cohort.

**Supplementary Table S4:** Comparison of AUCs across different omics datatype combinations for different subtypes in the breast cancer cohort.

**Supplementary Table S5:** Drug-response datasets summary.

## References

1. Menyhart, O. and Gyorffy, B. (2021) Multi-omics approaches in cancer research with applications in tumor subtyping, prognosis, and diagnosis. Comput Struct Biotechnol J, 19, 949–960.

2. Heo, Y.J., Hwa, C., Lee, G.H., Park, J.M. and An, J.Y. (2021) Integrative Multi-Omics Approaches in Cancer Research: From Biological Networks to Clinical Subtypes. Mol Cells, 44, 433–443.

3. Nicora, G., Vitali, F., Dagliati, A., Geifman, N. and Bellazzi, R. (2020) Integrated Multi-Omics Analyses in Oncology: A Review of Machine Learning Methods and Tools. Front Oncol, 10, 1030.

4. Lu, M. and Zhan, X. (2018) The crucial role of multiomic approach in cancer research and clinically relevant outcomes. EPMA J, 9, 77–102.

5. Tran, K.A., Kondrashova, O., Bradley, A., Williams, E.D., Pearson, J.V. and Waddell, N. (2021) Deep learning in cancer diagnosis, prognosis and treatment selection. Genome Med, 13, 152.

6. Wekesa, J.S. and Kimwele, M. (2023) A review of multi-omics data integration through deep learning approaches for disease diagnosis, prognosis, and treatment. Front Genet, 14, 1199087.

7. Chaudhary, K., Poirion, O.B., Lu, L. and Garmire, L.X. (2018) Deep Learning-Based Multi-Omics Integration Robustly Predicts Survival in Liver Cancer. Clin Cancer Res, 24, 1248–1259.

8. Lin, E., Lin, C.H. and Lane, H.Y. (2021) Deep Learning with Neuroimaging and Genomics in Alzheimer’s Disease. Int J Mol Sci, 22.

9. Albaradei, S., Thafar, M., Alsaedi, A., Van Neste, C., Gojobori, T., Essack, M. and Gao, X. (2021) Machine learning and deep learning methods that use omics data for metastasis prediction. Comput Struct Biotechnol J, 19, 5008–5018.

10. Poirion, O.B., Jing, Z., Chaudhary, K., Huang, S. and Garmire, L.X. (2021) DeepProg: an ensemble of deep-learning and machine-learning models for prognosis prediction using multi-omics data. Genome Med, 13, 112.

11. Chai, H., Zhou, X., Zhang, Z., Rao, J., Zhao, H. and Yang, Y. (2021) Integrating multi-omics data through deep learning for accurate cancer prognosis prediction. Comput Biol Med, 134, 104481.

12. Xie, G., Dong, C., Kong, Y., Zhong, J.F., Li, M. and Wang, K. (2019) Group Lasso Regularized Deep Learning for Cancer Prognosis from Multi-Omics and Clinical Features. Genes (Basel), 10.

13. Sharifi-Noghabi, H., Zolotareva, O., Collins, C.C. and Ester, M. (2019) MOLI: multi-omics late integration with deep neural networks for drug response prediction. Bioinformatics, 35, i501–i509.

14. Simidjievski, N., Bodnar, C., Tariq, I., Scherer, P., Andres Terre, H., Shams, Z., Jamnik, M. and Lio, P. (2019) Variational Autoencoders for Cancer Data Integration: Design Principles and Computational Practice. Front Genet, 10, 1205.

15. Wang, T., Shao, W., Huang, Z., Tang, H., Zhang, J., Ding, Z. and Huang, K. (2021) MOGONET integrates multi-omics data using graph convolutional networks allowing patient classification and biomarker identification. Nat Commun, 12, 3445.

16. Benkirane, H., Pradat, Y., Michiels, S. and Cournede, P.H. (2023) CustOmics: A versatile deep-learning based strategy for multi-omics integration. PLoS Comput Biol, 19, e1010921.

17. Liu, B., Wei, Y., Zhang, Y. and Yang, Q. (2017), Proceedings of the Twenty-Sixth International Joint Conference on Artificial Intelligence*, {*IJCAI*-*17*}*, pp. 2287--2293.

18. Kang, M., Ko, E. and Mersha, T.B. (2022) A roadmap for multi-omics data integration using deep learning. Brief Bioinform, 23.

19. Sharma, A., Vans, E., Shigemizu, D., Boroevich, K.A. and Tsunoda, T. (2019) DeepInsight: A methodology to transform a non-image data to an image for convolution neural network architecture. Sci Rep, 9, 11399.

20. Bazgir, O., Zhang, R., Dhruba, S.R., Rahman, R., Ghosh, S. and Pal, R. (2020) Representation of features as images with neighborhood dependencies for compatibility with convolutional neural networks. Nat Commun, 11, 4391.

21. Zhu, Y., Brettin, T., Xia, F., Partin, A., Shukla, M., Yoo, H., Evrard, Y.A., Doroshow, J.H. and Stevens, R.L. (2021) Converting tabular data into images for deep learning with convolutional neural networks. Sci Rep, 11, 11325.

22. Sharma, A., Lysenko, A., Boroevich, K.A. and Tsunoda, T. (2023) DeepInsight-3D architecture for anti-cancer drug response prediction with deep-learning on multi-omics. Sci Rep, 13, 2483.

23. Krzywinski, M., Schein, J., Birol, I., Connors, J., Gascoyne, R., Horsman, D., Jones, S.J. and Marra, M.A. (2009) Circos: an information aesthetic for comparative genomics. Genome Res, 19, 1639–1645.

24. Deng, J., Dong, W., Socher, R., Li, L.J., Kai, L. and Li, F.-F. (2009), 2009 IEEE Conference on Computer Vision and Pattern Recognition, pp. 248-255.

25. Gneiting, T. and Raftery, A.E. (2007) Strictly Proper Scoring Rules, Prediction, and Estimation. Journal of the American Statistical Association, 102, 359–378.

26. Khan, H.A. (2018) DM-L Based Feature Extraction and Classifier Ensemble for Object Recognition. Journal of Signal and Information Processing, Vol.09 No.02, 19.

27. Erickson, N., Mueller, J., Shirkov, A., Zhang, H., Larroy, P., Li, M. and Smola, A. (2020) AutoGluon-Tabular: Robust and Accurate AutoML for Structured Data. arXiv e-prints, arXiv:2003.06505.

28. Lin, T.-Y., RoyChowdhury, A. and Maji, S. (2015) Bilinear CNNs for Fine-grained Visual Recognition. arXiv e-prints, arXiv:1504.07889.

29. Huang, G., Liu, Z., van der Maaten, L. and Weinberger, K.Q. (2016) Densely Connected Convolutional Networks. arXiv e-prints, arXiv:1608.06993.

30. Tan, M. and Le, Q.V. (2021) EfficientNetV2: Smaller Models and Faster Training. *arXiv e-prints*, arXiv:2104.00298.

31. Al-Sabaawi, A., Ibrahim, H.M., Arkah, Z.M., Al-Amidie, M. and Alzubaidi, L. (2021) In Abraham, A., Piuri, V., Gandhi, N., Siarry, P., Kaklauskas, A. and Madureira, A. (eds.), Intelligent Systems Design and Applications. Springer International Publishing, Cham, pp. 171–180.

32. Lundberg, S. and Lee, S.-I. (2017) A Unified Approach to Interpreting Model Predictions. arXiv e-prints, arXiv:1705.07874.

33. Goldman, M.J., Craft, B., Hastie, M., Repecka, K., McDade, F., Kamath, A., Banerjee, A., Luo, Y., Rogers, D., Brooks, A.N. et al. (2020) Visualizing and interpreting cancer genomics data via the Xena platform. Nat Biotechnol, 38, 675–678.

34. Colaprico, A., Silva, T.C., Olsen, C., Garofano, L., Cava, C., Garolini, D., Sabedot, T.S., Malta, T.M., Pagnotta, S.M., Castiglioni, I. et al. (2016) TCGAbiolinks: an R/Bioconductor package for integrative analysis of TCGA data. Nucleic Acids Res, 44, e71.

35. Ciriello, G., Gatza, M.L., Beck, A.H., Wilkerson, M.D., Rhie, S.K., Pastore, A., Zhang, H., McLellan, M., Yau, C., Kandoth, C. et al. (2015) Comprehensive Molecular Portraits of Invasive Lobular Breast Cancer. Cell, 163, 506–519.

36. Kim, H. and Park, H. (2007) Sparse non-negative matrix factorizations via alternating non-negativity-constrained least squares for microarray data analysis. Bioinformatics, 23, 1495–1502.

37. Kolberg, L., Raudvere, U., Kuzmin, I., Vilo, J. and Peterson, H. (2020) gprofiler2--an R package for gene list functional enrichment analysis and namespace conversion toolset g:Profiler. F1000Res, 9.

38. Kramer, A., Green, J., Pollard, J., Jr. and Tugendreich, S. (2014) Causal analysis approaches in Ingenuity Pathway Analysis. Bioinformatics, 30, 523–530.

39. Gu, Z., Gu, L., Eils, R., Schlesner, M. and Brors, B. (2014) circlize Implements and enhances circular visualization in R. Bioinformatics, 30, 2811–2812.

40. Lundberg, S.M. and Lee, S.-I. (2017) In Guyon, I., Luxburg, U. V., Bengio, S., Wallach, H., Fergus, R., Vishwanathan, S. and Garnett, R. (eds.), Vol. 30.

41. Gao, C., Wei, J., Tang, T. and Huang, Z. (2020) Role of microRNA-33a in malignant cells. Oncol Lett, 20, 2537–2556.

42. Bastien, R.R., Rodriguez-Lescure, A., Ebbert, M.T., Prat, A., Munarriz, B., Rowe, L., Miller, P., Ruiz-Borrego, M., Anderson, D., Lyons, B. et al. (2012) PAM50 breast cancer subtyping by RT-qPCR and concordance with standard clinical molecular markers. BMC Med Genomics, 5, 44.

43. Prat, A., Parker, J.S., Fan, C. and Perou, C.M. (2012) PAM50 assay and the three-gene model for identifying the major and clinically relevant molecular subtypes of breast cancer. Breast Cancer Res Treat, 135, 301–306.

44. Prat, A. and Perou, C.M. (2009) Mammary development meets cancer genomics. Nat Med, 15, 842–844.

45. Korkola, J.E., DeVries, S., Fridlyand, J., Hwang, E.S., Estep, A.L., Chen, Y.Y., Chew, K.L., Dairkee, S.H., Jensen, R.M. and Waldman, F.M. (2003) Differentiation of lobular versus ductal breast carcinomas by expression microarray analysis. Cancer Res, 63, 7167–7175.

