## Supplementary figures for "OmicsFootPrint: a framework to integrate and interpret multi-omics data using circular images and deep neural networks"

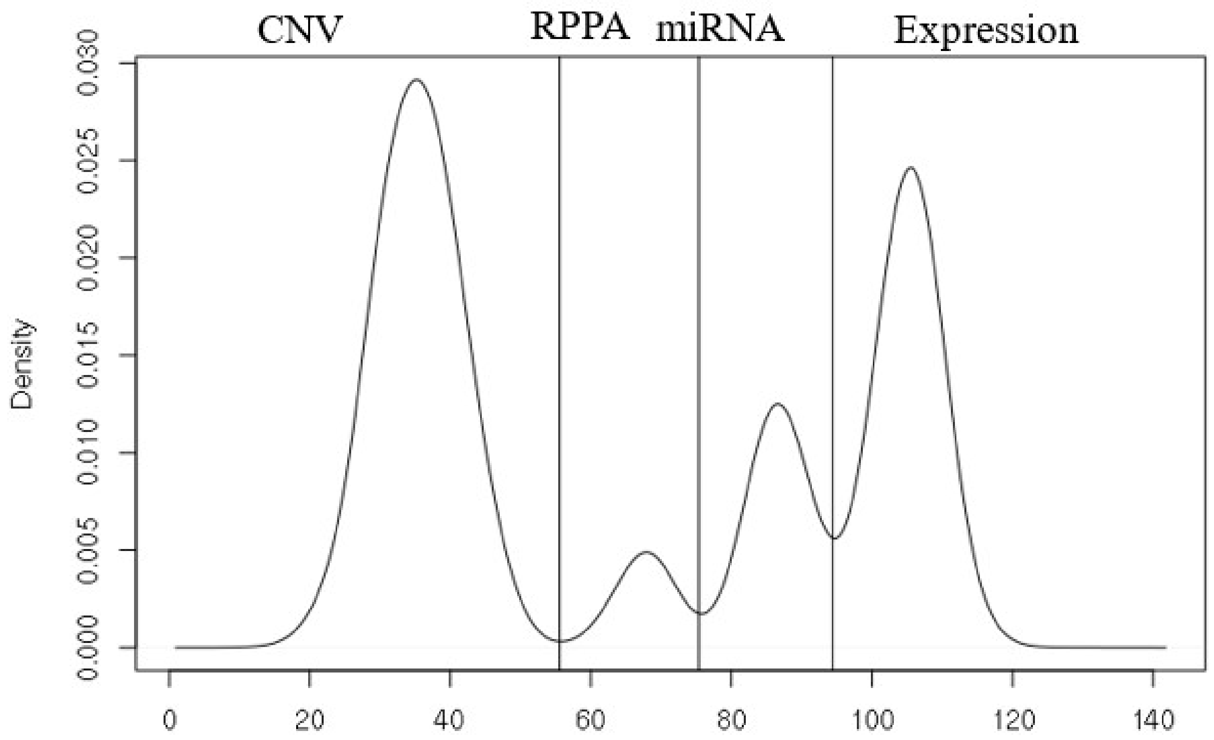


**Supplementary Figure S1:** Distribution of the distances of peaks to the center of the image. The density curve shows that peaks are enriched by data type. This property is used to determine the omics data type of any peak point on the circular image in a data-driven way.
