## Supplementary tables for "OmicsFootPrint: a framework to integrate and interpret multi-omics data using circular images and deep neural networks"

Table S1: Drug-response datasets summary

| Drug | Training Set Source | Test Set Source | # of Training samples | # of Test Samples |
|---|---|---|---|---|
| **Paclitaxel** | GDSC | PDX | 389 (R:363, S:26) | 43 (R:38, S:5) |
| **Gemcitabine** | GDSC | PDX | 844 (R:790, S:54) | 25 (R:18, S:7) |
| **Cetuximab** | GDSC | PDX | 856 (R:735, S:121) | 60 (R:55, S:5) |
| **Erlotinib** | GDSC | PDX | 362 (R:298, S:64) | 21 (R:18, S:3) |
| **Docetaxel** | GDSC | TCGA | 829 (R:764, S:65) | 16 (R:8, S:8) |
| **Cisplatin** | GDSC | TCGA | 829 (R:752, S:77) | 66 (R:6, S:60) |
| **Gemcitabine** | GDSC | TCGA | 844 (R:790, S:54) | 57 (R:36, S:21) |

### Table S2: Comparison of AUCs across different architectures for different subtypes in the lung cohort

| DNN Architectures | Seed_1001 | Seed_1002 | Seed_1003 | Seed_1004 | Seed_1005 | Seed_1006 | Seed_1007 | Seed_1008 | Seed_1009 | Seed_1010 | Mean | Median | SD | SE | IQR |
|---|---|---|---|---|---|---|---|---|---|---|---|---|---|---|---|
| EfficientNetV2 | 0.99 | 0.97 | 1 | 0.95 | 0.95 | 0.99 | 0.98 | 0.97 | 0.98 | 0.97 | **0.98** | **0.98** | 0.02 | 0.01 | 0.02 |
| DenseNet121 | 1 | 0.96 | 0.99 | 0.93 | 0.95 | 0.98 | 0.97 | 0.97 | 0.99 | 0.97 | 0.97 | 0.97 | 0.02 | 0.01 | 0.03 |
| Bilinear | 1 | 0.96 | 0.99 | 0.96 | 0.97 | 0.98 | 0.96 | 0.97 | 0.99 | 0.96 | 0.97 | 0.97 | 0.02 | 0 | 0.03 |
| VGG16.AutoGluon | 0.84 | 0.9 | 0.91 | 0.9 | 0.93 | 0.93 | 0.87 | 0.94 | 0.87 | 0.87 | 0.90 | 0.90 | 0.03 | 0.01 | 0.06 |

### Table S3: Comparison of AUCs across different omics combination for different subtypes in the breast cancer cohort

| DNN Architectures | Subtype | Seed_1001 | Seed_1002 | Seed_1003 | Seed_1004 | Seed_1005 | Seed_1006 | Seed_1007 | Seed_1008 | Seed_1009 | Seed_1010 | Mean | Median | SD | SE | IQR |
|---|---|---|---|---|---|---|---|---|---|---|---|---|---|---|---|---|
| EfficientNetV2 | Basal | 0.97 | 0.98 | 0.98 | 0.99 | 0.96 | 0.97 | 0.94 | 0.94 | 0.98 | 1 | **0.97** | **0.98** | 0.02 | 0.01 | 0.02 |
| EfficientNetV2 | HER2 | 0.96 | 0.99 | 0.88 | 0.77 | 0.84 | 0.93 | 0.79 | 0.93 | 0.89 | 0.89 | **0.89** | **0.89** | 0.07 | 0.02 | 0.08 |
| EfficientNetV2 | LumA | 0.81 | 0.77 | 0.82 | 0.86 | 0.85 | 0.81 | 0.77 | 0.77 | 0.82 | 0.82 | **0.81** | **0.82** | 0.03 | 0.01 | 0.04 |
| EfficientNetV2 | LumB | 0.67 | 0.74 | 0.69 | 0.81 | 0.8 | 0.76 | 0.73 | 0.74 | 0.75 | 0.74 | **0.74** | **0.74** | 0.04 | 0.01 | 0.03 |
| Bilinear | Basal | 0.98 | 0.99 | 0.95 | 1 | 0.89 | 0.98 | 1 | 0.93 | 1 | 0.97 | **0.97** | **0.98** | 0.04 | 0.01 | 0.04 |
| Bilinear | HER2 | 0.92 | 0.84 | 0.81 | 0.89 | 0.91 | 0.78 | 0.89 | 0.73 | 0.94 | 0.93 | 0.86 | **0.89** | 0.07 | 0.02 | 0.1 |
| Bilinear | LumA | 0.84 | 0.78 | 0.74 | 0.77 | 0.77 | 0.79 | 0.81 | 0.78 | 0.83 | 0.77 | 0.79 | 0.78 | 0.03 | 0.01 | 0.04 |
| Bilinear | LumB | 0.63 | 0.61 | 0.51 | 0.56 | 0.49 | 0.57 | 0.68 | 0.59 | 0.68 | 0.54 | 0.59 | 0.58 | 0.07 | 0.02 | 0.08 |
| DenseNet121 | Basal | 0.91 | 0.97 | 0.89 | 0.98 | 0.96 | 0.99 | 0.96 | 0.9 | 0.95 | 0.95 | 0.95 | 0.96 | 0.03 | 0.01 | 0.05 |
| DenseNet121 | HER2 | 0.72 | 0.85 | 0.7 | 0.88 | 0.95 | 0.94 | 0.75 | 0.84 | 0.89 | 0.8 | 0.83 | 0.85 | 0.09 | 0.03 | 0.12 |
| DenseNet121 | LumA | 0.83 | 0.79 | 0.73 | 0.84 | 0.82 | 0.77 | 0.79 | 0.77 | 0.79 | 0.77 | 0.79 | 0.79 | 0.03 | 0.01 | 0.04 |
| DenseNet121 | LumB | 0.65 | 0.8 | 0.74 | 0.67 | 0.76 | 0.77 | 0.67 | 0.53 | 0.75 | 0.62 | 0.70 | 0.71 | 0.08 | 0.03 | 0.1 |
| VGG16.AutoGluon | Basal | 0.98 | 0.94 | 0.95 | 0.95 | 0.92 | 0.87 | 0.93 | 0.8 | 0.95 | 0.92 | 0.92 | 0.94 | 0.05 | 0.02 | 0.03 |
| VGG16.AutoGluon | HER2 | 0.85 | 0.7 | 0.64 | 0.64 | 0.74 | 0.63 | 0.55 | 0.78 | 0.73 | 0.63 | 0.69 | 0.67 | 0.09 | 0.03 | 0.1 |
| VGG16.AutoGluon | LumA | 0.84 | 0.75 | 0.82 | 0.8 | 0.76 | 0.78 | 0.8 | 0.78 | 0.81 | 0.8 | 0.79 | 0.80 | 0.03 | 0.01 | 0.03 |
| VGG16.AutoGluon | LumB | 0.75 | 0.58 | 0.69 | 0.65 | 0.7 | 0.74 | 0.68 | 0.76 | 0.6 | 0.68 | 0.68 | 0.69 | 0.06 | 0.02 | 0.07 |

### Table S4: Comparison of AUCs across different omics datatype combinations for different subtypes in the lung cohort

| Omics Combination | Seed_1001 | Seed_1002 | Seed_1003 | Seed_1004 | Seed_1005 | Seed_1006 | Seed_1007 | Seed_1008 | Seed_1009 | Seed_1010 | Mean | Median | SD | SE | IQR |
|---|---|---|---|---|---|---|---|---|---|---|---|---|---|---|---|
| expr_only | 0.95 | 0.89 | 0.9 | 0.88 | 0.92 | 0.97 | 0.92 | 0.9 | 0.98 | 0.92 | 0.92 | 0.92 | 0.03 | 0.01 | 0.04 |
| expr_cnv | 0.91 | 0.9 | 0.9 | 0.89 | 0.93 | 0.91 | 0.88 | 0.97 | 0.91 | 0.92 | 0.91 | 0.91 | 0.02 | 0.01 | 0.02 |
| expr_cnv_rppa | 0.92 | 0.94 | 0.94 | 0.87 | 0.95 | 0.97 | 0.94 | 0.89 | 0.97 | 0.94 | **0.94** | **0.93** | 0.03 | 0.01 | 0.02 |

### Table S5: Comparison of AUCs across different omics datatype combinations for different subtypes in the breast cancer cohort

| Omics Combination | Subtype | Seed_1001 | Seed_1002 | Seed_1003 | Seed_1004 | Seed_1005 | Seed_1006 | Seed_1007 | Seed_1008 | Seed_1009 | Seed_1010 | Mean | Median | SD | SE | IQR | Overall Median |
|---|---|---|---|---|---|---|---|---|---|---|---|---|---|---|---|---|---|
| expr_cnv_rppa | Basal | 0.95 | 0.97 | 0.98 | 0.98 | 0.97 | 1 | 0.95 | 0.92 | 0.97 | 0.98 | **0.97** | 0.97 | 0.02 | 0.007 | 0.025 | |
| expr_cnv_rppa | HER2 | 0.89 | 0.89 | 0.74 | 0.73 | 0.94 | 0.89 | 0.84 | 0.68 | 0.89 | 0.89 | 0.84 | **0.89** | 0.09 | 0.028 | 0.125 | |
| expr_cnv_rppa | LumA | 0.92 | 0.88 | 0.82 | 0.86 | 0.85 | 0.84 | 0.84 | 0.83 | 0.86 | 0.86 | **0.86** | **0.86** | 0.03 | 0.009 | 0.02 | |
| expr_cnv_rppa | LumB | 0.89 | 0.78 | 0.74 | 0.77 | 0.7 | 0.72 | 0.77 | 0.76 | 0.79 | 0.72 | **0.76** | **0.77** | 0.05 | 0.017 | 0.052 | **0.86** |
| expr_cnv | Basal | 0.93 | 0.97 | 0.99 | 0.99 | 0.98 | 0.98 | 0.99 | 0.94 | 0.97 | 0.98 | **0.97** | **0.98** | 0.02 | 0.007 | 0.017 | |
| expr_cnv | HER2 | 0.92 | 0.86 | 0.74 | 0.77 | 0.99 | 0.84 | 0.89 | 0.87 | 0.84 | 0.7 | 0.84 | 0.85 | 0.09 | 0.027 | 0.098 | |
| expr_cnv | LumA | 0.89 | 0.73 | 0.83 | 0.84 | 0.77 | 0.88 | 0.86 | 0.83 | 0.85 | 0.73 | 0.82 | 0.84 | 0.06 | 0.018 | 0.073 | |
| expr_cnv | LumB | 0.81 | 0.67 | 0.73 | 0.7 | 0.74 | 0.81 | 0.72 | 0.71 | 0.76 | 0.6 | 0.73 | 0.73 | 0.06 | 0.02 | 0.053 | 0.84 |
| expr_only | Basal | 0.97 | 0.98 | 0.97 | 0.99 | 0.97 | 0.98 | 0.95 | 0.95 | 0.95 | 0.99 | 0.97 | 0.97 | 0.02 | 0.005 | 0.025 | |
| expr_only | HER2 | 0.93 | 0.79 | 0.83 | 0.9 | 0.78 | 0.89 | 0.84 | 0.74 | 0.99 | 0.89 | **0.86** | 0.87 | 0.08 | 0.024 | 0.098 | |
| expr_only | LumA | 0.87 | 0.8 | 0.83 | 0.85 | 0.89 | 0.86 | 0.85 | 0.78 | 0.84 | 0.84 | 0.84 | 0.85 | 0.03 | 0.01 | 0.025 | |
| expr_only | LumB | 0.67 | 0.66 | 0.73 | 0.78 | 0.8 | 0.77 | 0.78 | 0.68 | 0.78 | 0.73 | 0.74 | 0.75 | 0.05 | 0.016 | 0.088 | 0.845 |
| cnv_only | Basal | 0.73 | 0.76 | 0.73 | 0.71 | 0.71 | 0.83 | 0.7 | 0.82 | 0.84 | 0.71 | 0.75 | 0.73 | 0.06 | 0.017 | 0.095 | |
| cnv_only | HER2 | 0.78 | 0.57 | 0.55 | 0.68 | 0.66 | 0.71 | 0.67 | 0.57 | 0.6 | 0.7 | 0.65 | 0.67 | 0.07 | 0.023 | 0.117 | |
| cnv_only | LumA | 0.76 | 0.67 | 0.78 | 0.77 | 0.67 | 0.74 | 0.74 | 0.78 | 0.77 | 0.82 | 0.75 | 0.77 | 0.05 | 0.015 | 0.037 | |
| cnv_only | LumB | 0.69 | 0.63 | 0.64 | 0.58 | 0.52 | 0.6 | 0.59 | 0.61 | 0.67 | 0.56 | 0.61 | 0.61 | 0.05 | 0.016 | 0.055 | 0.7 |
